# Antibiotic breakdown by susceptible bacteria enhances the establishment of β-lactam resistant mutants

**DOI:** 10.1101/2021.04.20.440616

**Authors:** Manja Saebelfeld, Suman G. Das, Jorn Brink, Arno Hagenbeek, Joachim Krug, J. Arjan G.M. de Visser

## Abstract

For a better understanding of the evolution of antibiotic resistance, it is imperative to study the factors that determine the initial establishment of mutant resistance alleles. In addition to the antibiotic concentration, the establishment of resistance alleles may be affected by interactions with the surrounding susceptible cells from which they derive, for instance via the release of nutrients or removal of the antibiotic. Here, we investigate the effects of social interactions with surrounding susceptible cells on the establishment of *Escherichia coli* mutants with increasing β-lactamase activity (i.e. the capacity to hydrolyze β-lactam antibiotics) from single cells under the exposure of the antibiotic cefotaxime on agar plates. We find that mutant establishment probability is increased in the presence of susceptible cells due to the active breakdown of the antibiotic, but the rate of breakdown by the susceptible strain is much higher than expected based on its low enzymatic activity. A detailed theoretical model suggests that this observation can be explained by cell filamentation causing delayed lysis. While susceptible cells may hamper the spread of higher-resistant β-lactamase mutants at relatively high frequencies, our findings show that they could promote establishment during their emergence.

## 1 Introduction

Antibiotic resistance has become a worldwide concern, causing 700,000 deaths annually due to failed treatments (IACG, 2019). As the development of novel drugs does not appear to be promising in the long-term, other strategies of slowing or halting the emergence of antibiotic resistance have been explored, including diverse prevention strategies (e.g. rational use of antibiotics and infection control), the use of biologics or adjuvants to disturb quorum sensing of resistant bacteria, the use of bacteriophages (reviewed in Aslam et al., 2018) and the exploitation of the evolutionary potential and interactions of resistance to develop new treatment strategies (Schenk et al., 2012; Palmer and Kishony, 2013; Baym et al., 2016; Furusawa et al., 2018).

For a better understanding of the evolution of antibiotic resistance, it is not only important to study the spread of resistant bacteria, but also the factors that determine the emergence of new resistance alleles in the first place (Alexander and MacLean, 2020). Resistance alleles can be acquired via mutation or horizontal gene transfer (Palmer and Kishony, 2013; Blair et al., 2015). In either case, the resistant allele is initially present at a very low frequency and thus prone to random extinction. Population genetics theory predicts that only when an allele has survived extinction by genetic drift and reaches an absolute frequency roughly equal to the inverse of its selection coefficient, its course becomes dominated by selection in an environment where it is beneficial (Haldane, 1927; Patwa and Wahl, 2008). Mutations that have survived genetic drift are often called “established” (Levy et al., 2015; Alexander and MacLean, 2020), although they may still be driven to extinction by the competition with other mutants (Gerrish and Lenski, 1998; Rozen et al., 2002). The few existing empirical studies on the establishment of rare alleles have shown that this stochastic process of “drift loss” is influenced by the initial allele frequency and the fitness benefit of the allele (Chelo et al., 2013; Gifford et al., 2013).

Only a handful of studies so far have systematically investigated the establishment of single bacterial cells expressing an antibiotic-resistant allele. Those studies found that the probability of a resistant mutant to establish depends on the type and concentration of nutrients (Coates et al., 2018) and antibiotics (Coates et al., 2018; Alexander and MacLean, 2020). Alexander and MacLean (2020) found that the establishment probability of a single streptomycin-resistant *Pseudomonas aeruginosa* cell was only 5% at a concentration as low as 1/8 of the strains’ minimum inhibitory concentration (MIC) in the absence of a wildtype population. Interestingly, when the same mutant was introduced into a large population of wildtype cells, its establishment was strongly increased under the same streptomycin concentrations compared to the absence of the wildtype. The authors speculated that this effect was due to the removal of the antibiotic from the environment via binding to wildtype cell components (Alexander and MacLean, 2020).

Determining factors that influence the establishment of antibiotic-resistance alleles could have implications in a clinical context, for instance by adjusting drug dosing strategies to a point where the establishment of *de novo* mutations is strongly reduced (Alexander and MacLean, 2020). For this, interactions between newly arising resistance alleles and nearby sensitive cells must be understood better. Apart from drug removal by binding to cellular components, other common goods, such as enzymes causing antibiotic hydrolysis or the release of nutrients from killed cells, may play a role. A good model system to study such social interactions is TEM-1 β-lactamase. This enzyme is expressed in the periplasmic space of the bacterial cell wall, where it hydrolyses β-lactam antibiotics such as penicillins. This leads to a reduction of the antibiotic concentration in the environment and may support the growth of susceptible cells (Brown et al., 2009; Medaney et al., 2016; Nicoloff and Andersson, 2016). Such cooperative behavior is expected to be more pronounced in structured environments, such as biofilms, than in unstructured ones, due to stronger effects of enzymatic breakdown on local antibiotic concentrations (Medaney et al., 2016; Rojo-Molinero et al., 2019). If and to what extent such interactions affect the establishment of β-lactamase mutants conferring higher resistance, is unknown.

Here, we tested the effect of relatively antibiotic-susceptible cells on the establishment of higher-resistance TEM-1 β-lactamase alleles. In particular, we distinguished between possible effects of increased nutrient availability due to cell lysis and drug removal via β-lactamase activity or binding to cellular components. We used the β-lactam antibiotic cefotaxime (CTX) and different strains of the bacterium *Escherichia coli*, expressing β-lactamase alleles with varying activity against CTX. We simulated the establishment process of CTX-resistant *E. coli* by introducing a low number of mutant cells into populations of more susceptible cells on CTX-containing agar plates. Our results revealed substantial increases in mutant establishment probability by susceptible cells. By using heat-killed susceptible cells and measuring the CTX-reducing capacity of mutants compared to the wildtype, we distinguished between the effects from nutrient addition by cell lysis and antibiotic removal via either CTX binding or hydrolysis on mutant establishment. Lastly, to understand an unexpectedly large effect on mutant establishment by the ancestral TEM-1 strain, we used a detailed dynamical model. This model suggests that certain population dynamic parameters, such as cell filamentation and β-lactamase synthesis rate, may have played a crucial role during the establishment of the higher-resistance mutants.

## 2 Material and Methods

### 2.1 Strains and culture conditions

Several strains, derived from *Escherichia coli* strain MG1655 were used for the experiments. This strain had previously been modified with either a YFP (yellow fluorescent protein) or a BFP (blue fluorescent protein) chromosomal marker cassette, containing a resistance gene for chloramphenicol (Gullberg et al., 2014). For the current study, the chloramphenicol resistance was removed to serve as the Wildtype strain (Table 1). Into each of the two fluorescently-marked Wildtype strains, each of four TEM variants was inserted into the *galK* locus: TEM-1 (from now on referred to as “Ancestral” strain), and three mutant alleles with either 1, 2 or 3 point mutations in the TEM gene (from now on referred to as “Single mutant”, “Double mutant” and “Triple mutant”, respectively; cf. Table 1). All TEM loci are under the control of the *lacI* repressor and are expressed by adding 50 μM Isopropyl β-D-1-thiogalactopyranoside (IPTG) to the growth medium. While the TEM-1 allele confers very low activity towards CTX, the three mutants show increasing CTX resistance with each additional substitution. The minimum inhibitory concentration (MIC) for CTX was determined per strain in duplicates (for each of the YFP and BFP variants), using 2-fold increases in CTX concentration in microtiter plates filled with 200 μl M9 minimal medium (containing 0.4% glucose, 0.2% casaminoacids, 2 μg/ml uracil and 1 μg/ml thiamine) and 50 μM IPTG, inoculated with 10^5^ cells and incubated for 24h at 37°C (cf. Table 1).

**Table 1:**
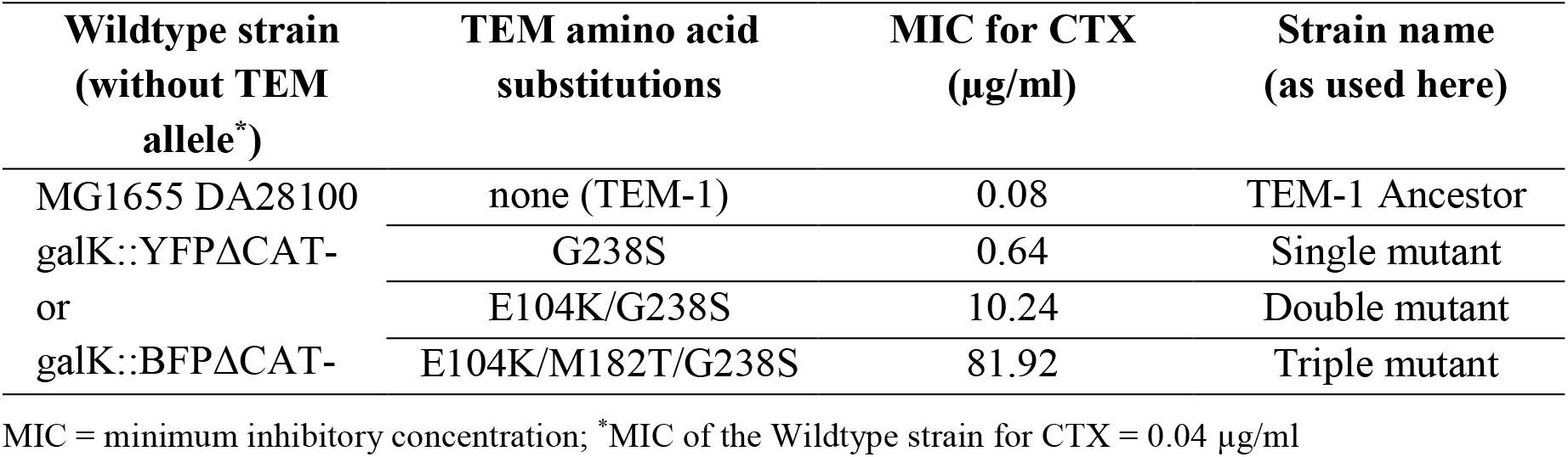
Overview of the used strains.

For setting up the experiments, all strains were grown overnight at 37°C and 250 rpm in 1-2 ml M9 medium. The cultures were either directly inoculated from the −80°C glycerol stocks or first streaked out on LB (Luria-Bertani) agar plates from which a single colony was picked into the liquid M9 medium. After overnight incubation, the cultures were serially diluted with phosphate-buffered saline (PBS) to the density needed for the particular experiment (see below). For the *Interaction experiments* on agar, the diluted cultures (or culture mixtures) were spread on 92 mm plates containing M9 medium (as above) with 1.5 % agar, 50 μM IPTG and the respective CTX concentration as specified by the particular experiment (see below).

### 2.2 Interaction experiment 1

First, we tested whether the previous observation that the establishment of an antibiotic-resistant mutant increases in the presence of susceptible cells (Alexander and MacLean, 2020), also applies to our TEM-1 β-lactamase system. For this, roughly 150 cells each of the three *E. coli* TEM mutants (Table 1) were introduced into populations of bacteria expressing the ancestral TEM-1 allele. All three mutants were plated alone or together with two densities of Ancestor cells under two CTX concentrations on M9 agar plates. The two tested CTX concentrations differed per mutant and were chosen to show single-cell establishment probabilities of about 50% and 80% compared to no CTX (Saebelfeld et al., bioRxiv preprint 2021). Overnight cultures of the mutants were serially diluted to approximately 3×10^3^ cells/ml, and the TEM-1 Ancestor overnight culture was serially diluted to 2×10^7^ cells/ml and 2×10^4^ cells/ml. The mutant dilutions were then mixed 1: 1 with either PBS or each of the two Ancestor density dilutions. Of those mixtures, 100 μl aliquots were spread on M9 plates containing the respective CTX concentrations (15 replicates per condition), using a bacterial spreader, resulting in ~150 cells of the respective mutant strain plated with either 0, ~10^3^, or ~10^6^ Ancestor cells. All used CTX concentrations were above the Ancestor MIC, allowing only for the mutant cells to grow into colonies. The plates were incubated at 37°C until the mutant colonies were big enough to count them unambiguously (20h-40h); the number of colony-forming units (CFUs) per plate was counted under white light, using a digital colony counter. To ensure that all counted colonies were indeed mutants, the BFP-expressing Ancestor strain, and the YFP-expressing mutant strains were used for this experiment. Within the lowest CTX concentration treatment for the Single mutant, one replicate of each tested Ancestor density was excluded from the dataset, as overlapping colonies prevented unambiguous CFU counts. After colony counting, all plates were checked under a fluorescence microscope (LEICA M165 FC), using GFP and CFP filters for detecting the YFP and BFP signals, respectively. For the Single mutant, a single colony containing the BFP Ancestor fluorophore was found within the 10^6^ ancestral cells treatment in each one replicate of the two tested CTX concentrations. These two colonies were subtracted from the CFU counts.

### 2.3 Interaction experiment 2

To test whether nutrient release from lysed cells and/or enzymatic breakdown of CTX could explain the observed positive effect on mutant establishment probability, we conducted a second plating experiment where approximately 150 Triple mutant cells were introduced into populations of ~10^6^ cells of either the TEM-1 Ancestor or the TEM Single mutant (with ~130-fold higher catalytic efficiency against CTX relative to TEM-1, see Salverda et al., 2011) under the same two CTX concentrations as in the *Interaction experiment 1*. In addition, the Triple mutant was introduced into populations of ~10^6^ cells of heat-killed Ancestor or heat-killed Single mutant cells, to test whether nutrient addition from cell lysis alone could explain the observed pattern.

To set up the experiment, half of the overnight culture of each of the YFP TEM-1 Ancestor and YFP Single mutant was heat-killed via incubation at 80°C for 3 hours and then put on ice for 20 minutes. An overnight culture of the BFP Triple mutant was serially diluted to 3×10^3^ cells/ml, and the alive and heat-killed overnight cultures of the YFP Ancestor and YFP Single mutant were diluted to 2×10^7^ cells/ml. The Triple mutant dilution was then mixed 1:1 with either PBS or the dilutions of alive or heat-killed Ancestor or alive Single mutant strains. 100 μl aliquots of these mixtures were then spread on M9 plates containing 1.44 or 1.76 μg/ml CTX (i.e. the same concentrations as used for the Triple mutant in the first interaction experiment; equivalent to 0.018 and 0.021% of the Triple mutant’s MIC), using 3 mm glass beads. As for the previous experiment, the used CTX concentrations were higher than the Ancestor and Single mutant MICs (see Table 1), so that only Triple mutant cells were expected to form colonies. The plates were incubated at 37°C until the colonies were big enough to count them unambiguously (16-20 h); the number of colony-forming units (CFUs) per plate was counted under white light, using a digital colony counter. To ensure that the counted colonies were Triple mutants, a fluorescence microscope was used as described above. No colonies expressing the YFP fluorophore (i.e. used for the Ancestor and Single mutant strains) were found. For each tested condition, 15 replicates were set up. To estimate the added number of cells per strain, the dilution containing only the Triple mutant was spread on 15 M9 plates without CTX. The alive Ancestor and alive Single mutant cell dilutions were further diluted to 10^3^ cells/ml; 200, 100 and 50 μl aliquots of these dilutions were spread on three LB agar plates each. All plates were incubated overnight at 37°C. Colony counts of the alive Ancestor and Single mutant cells showed that the initially added numbers in the experimental setup were comparable with 1.05×10^6^ and 1.01×10^6^ cells for each strain, respectively.

### 2.4 Bioassay

The antibiotic removal capacity of three strains was tested in a bioassay: the YFP TEM-1 Ancestor and TEM Single mutant, as used in the *Interaction experiment* 2, and in addition the YFP Wildtype strain (Table 1). This Wildtype strain does not contain a TEM allele and thus does not display any β-lactamase activity, but is otherwise genetically identical to both the TEM-1 Ancestor and Single mutant strains. This means that it contains the same number of penicillin-binding proteins (PBPs), which is the main target for CTX and thus controls for the difference of CTX removal via breakdown and target binding.

Overnight cultures of each strain were diluted 1:10 into 1 ml M9 medium, containing 50 μM IPTG and 1.76 μg/ml CTX (i.e. the highest tested concentration from the agar experiments) in three replicate tubes. In addition, heat-killed cells (as described above) of each strain were inoculated the same way. To control for spontaneous CTX breakdown under the tested conditions, three tubes were incubated in the absence of bacterial cells (CTX-only medium controls). For this, the M9 medium with CTX was mock-inoculated 1:10 with the M9 medium that was used for the overnight cultures the day before. All tubes were incubated for three hours at 37°C and 250 rpm. Although the CTX concentration was at least three-fold higher than the MIC of the strains, all alive cultures showed growth after the 3-hours incubation. This is likely to be attributed to an inoculum effect (Queenan et al., 2004), i.e. the increase of MIC with increasing inoculum size. The MIC is commonly tested at 5×10^5^ cells/ml while in our setup, the cultures contained densities of about 2×10^8^ cells/ml. We estimate the final density of the cultures to be ~10^9^ cells/ml after the 3-hours incubation.

After incubation, all tubes were centrifuged for 1 minute at 4000 rpm. The supernatant was sterilized through a 0.2 μm filter and used for setting up MIC assays. For this, the supernatant of each test tube was diluted in 1.35-fold steps with fresh M9 medium, containing 50 μM IPTG. In total, 11 putative CTX concentrations (i.e. assuming no removal) ranging from 0.04 to 0.79 μg/ml were tested per culture in 96-well plates. All wells were inoculated with ~10^5^ cells of the YFP TEM-1 Ancestor strain in a total volume of 200 μl per well. Growth of this strain in higher putative CTX concentrations would indicate the removal of CTX from the medium during the initial 3-hour culture incubation. Controls included (i) a filtration control: per culture, undiluted supernatant was mock-inoculated with M9_IPTG50_ medium, i.e. without bacterial cells to control for removal of alive cells after filtration; (ii) a growth control:M9_IPTG50_ medium without CTX was inoculated with ~10^5^ Ancestor cells; and (iii) a blank medium control consisting of M9_IPTG50_ medium only. The 96-well plates were incubated for 18 hours at 37°C (static). After incubation, OD at 600 nm of all wells was measured in a plate reader (Victor3™, PerkinElmer) without the plate lid. None of the supernatant or blank medium controls showed growth. The threshold to determine growth was set to an OD of 0.1. MICs were determined as the highest putative CTX concentration that did not show growth. To determine the amount of CTX removal from the initial cultures, the putative MIC per replicate was corrected for its dilution factor and the average MIC of the CTX-only medium controls. The resulting concentrations were then averaged across the three replicates per treatment.

### 2.5 Statistical analyses of the Interaction experiments

To test for a significant effect of the respective treatment on the number of mutant colony-forming units (CFUs), two-way ANOVAs were performed. For the analysis of *Interaction experiment 1*, the CFU counts were square-root transformed and the effects of CTX concentration and added number of Ancestor cells were included as fixed factors, followed by a Tukey’s HSD test. For the analysis of *Interaction experiment 2*, the effects of CTX concentration and background population were included as fixed factors after checking for homogeneous variances of the residuals and normal distribution of the data, followed by a Tukey’s HSD test. All statistical analyses were conducted in R v.3.6.2 (R Core Team, 2019) with the *car* package (Fox and Weisberg, 2019) to perform Levene tests; Figures 1 and 2 were produced using the package ggplot2 (Wickham, 2016).

**Figure 1:**
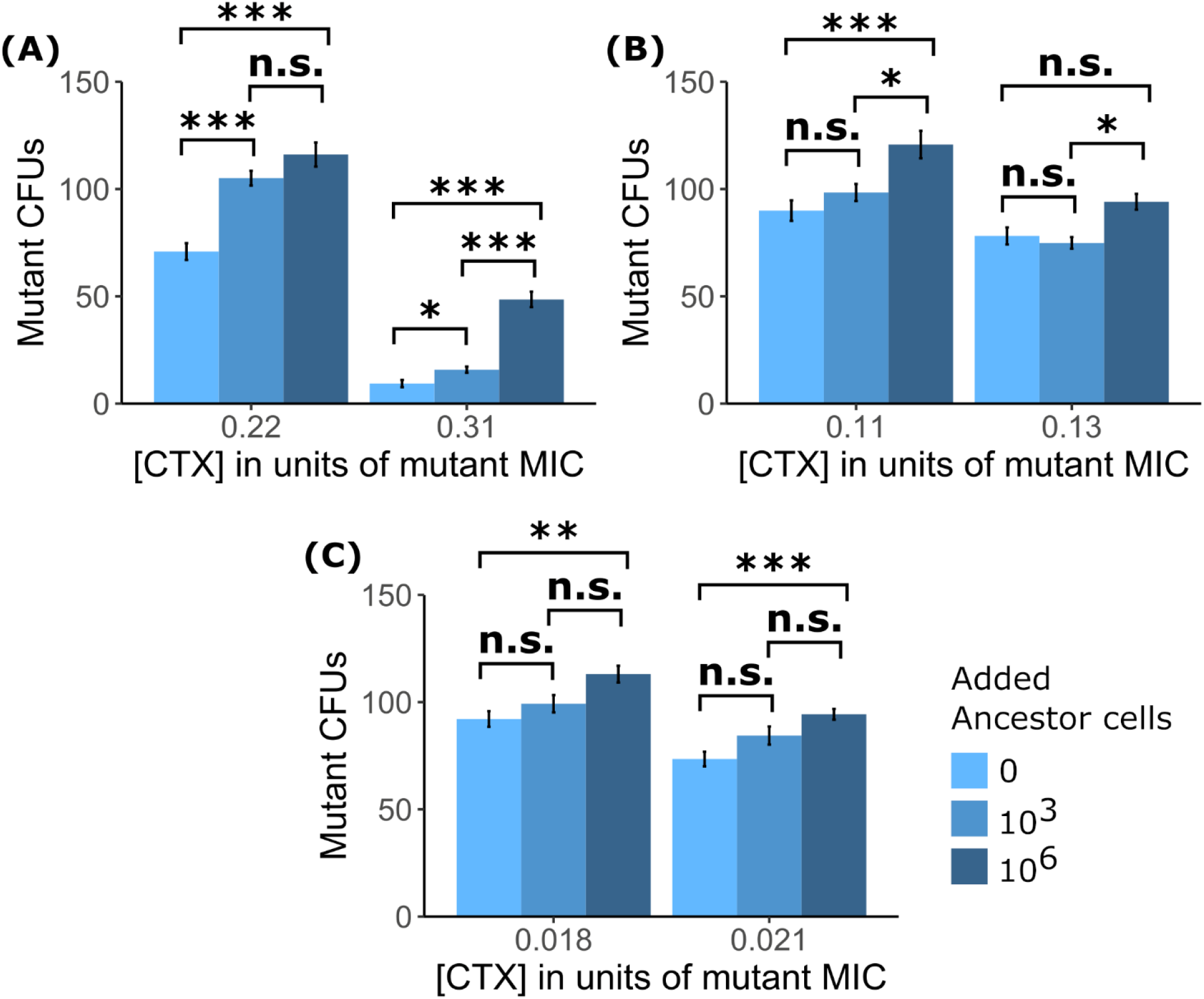
The establishment of single cells into visible colonies of the three tested TEM mutants is shown as counts of colony-forming units (CFUs) of **(A)** the Single mutant, **(B)** the Double mutant and **(C)** the Triple mutant under exposure to two CTX concentrations (shown as a fraction of each strain’s minimum inhibitory concentration – MIC) in either the absence of TEM-1 Ancestor cells or the presence of Ancestor cells at two different densities. Significance levels based on Tukey’s HSD tests are shown within CTX concentrations, between the number of added Ancestor cells after running two-way ANOVA’s over the whole dataset per mutant strain (cf. Supplement 1: Table S1). n.s. = not significant, *p<0.05, **p<0.01, ***p<0.001.

**Figure 2:**
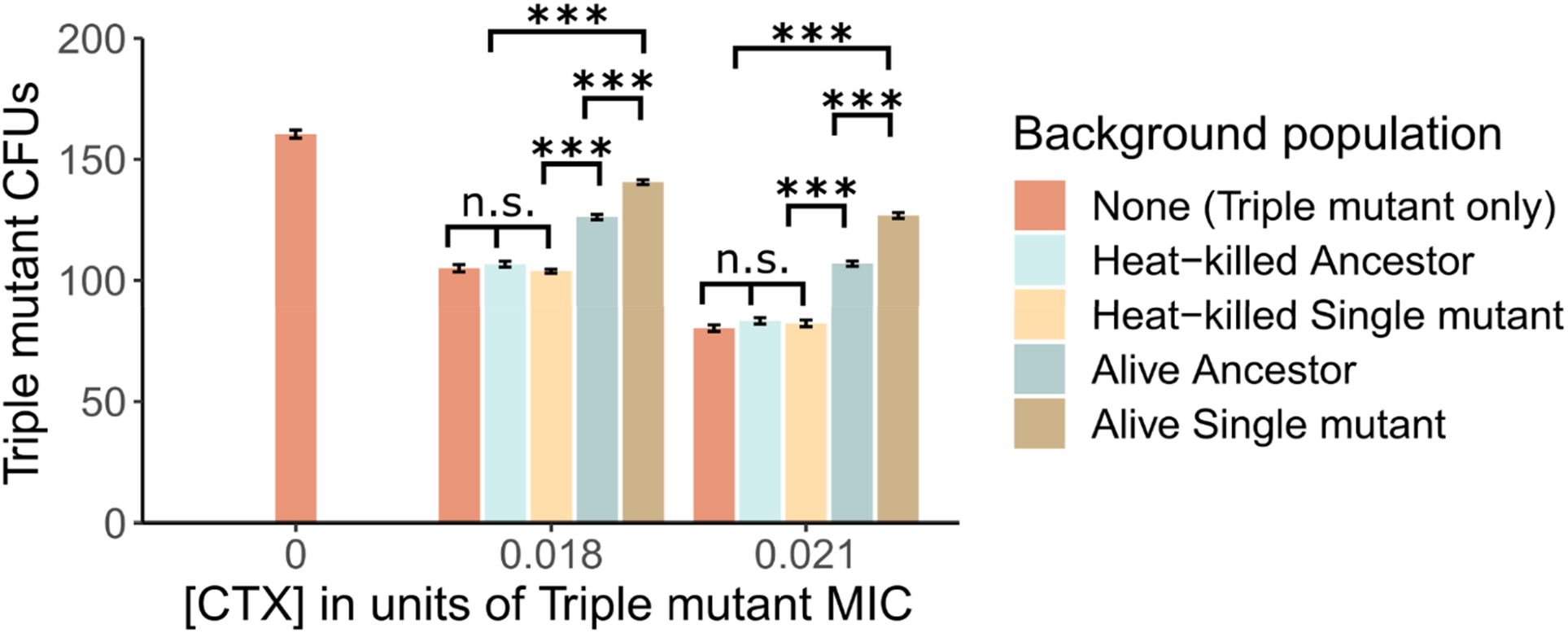
The establishment of Triple mutant cells is shown as counts of colony-forming units (CFUs) under the exposure of two CTX concentrations (shown as a fraction of the Triple mutant’s minimum inhibitory concentration – MIC), in the absence and presence of a “Background population”: 10^6^ alive or heat-killed TEM-1 Ancestor cells, and alive or heat-killed TEM Single mutant cells. In addition, the number of Triple mutant CFUs is shown in the absence of CTX. Significance levels based on Tukey’s HSD tests are shown within CTX concentrations, between the type of Background population after running two-way ANOVA’s over the whole dataset (cf. Supplement S2: Table S2). n.s. = not significant, ***p<0.001.

## 3 Results

### 3.1 Susceptible cells facilitate the establishment of β-lactamase mutants

In a first plating experiment, we tested whether the presence of TEM-1 Ancestor cells affected the establishment of three TEM mutants with increasing CTX resistance at two CTX concentrations. Generally, the presence of ancestral cells increased the number of established mutant cells of all three tested mutants (Figure 1), although this difference was not significant for all tested treatment comparisons (Supplement 1: Table S1). Moreover, establishment probability increased with Ancestral cell density under all conditions (Figure 1).

### 3.2 Mutant establishment is particularly affected by live cells expressing active β-lactamase

In a subsequent plating experiment, we tested whether nutrient access via lysed cells or CTX removal via enzymatic breakdown explains the positive effect of susceptible bacteria on the establishment of the resistant mutants. To do so, the Triple mutant was introduced into populations of either heat-killed or alive TEM-1 Ancestor or Single mutant cells under two CTX concentrations. When plated alone, the fraction of established Triple mutant cells, determined as colony-forming units (CFUs), decreased by approximately 34% and 50% in the low and high CTX concentrations compared to no CTX, respectively (Figure 2). The addition of heat-killed TEM-1 Ancestor or TEM Single mutant cells did not affect the establishment of the Triple mutant at either CTX concentration (Figure 2; Supplement 2: Table S2). However, the addition of alive Ancestor cells significantly increased the establishment of the Triple mutant under both CTX concentrations. This increase was further pronounced by the presence of alive Single mutant cells (Figure 2; Supplement 2: Table S2), suggesting an important role for enzymatic breakdown.

### 3.3 Test of CTX removal via target binding or breakdown

To confirm whether the increased establishment probability of the TEM Triple mutant in the presence of alive cells (Figure 2) is due to CTX removal, and examine the contribution of CTX binding and hydrolysis during this process, we performed a bioassay to estimate CTX removal for the Wildtype (no TEM), the TEM-1 Ancestor and the TEM Single mutant strains. We estimated the CTX removal rate of the three strains by culturing alive and heat-killed cells of each strain in medium with a supra-MIC CTX concentration for three hours and then using the cell-free medium in a standard MIC assay with the TEM-1 Ancestral strain (see Methods). Results show that the alive TEM-1 Ancestor and TEM Single mutant removed 0.29 and 0.43 μg/ml CTX (from the initial 1.79 μg/ml) from the liquid medium, respectively. Neither the Wildtype strain without the TEM allele nor the heat-killed mutants showed a decrease in CTX concentration (Table 2), supporting a significant role only for the β-lactamase in CTX removal.

**Table 2:** CTX removal from liquid medium in the bioassay by strains expressing different β-lactamase alleles or no β-lactamase, after 3 hours of incubation.

| Strain | Culture condition | CTX removal (in $\mu\text{g/ml}$) |
| --- | --- | --- |
| **TEM-1 Ancestor**<br>(low β-lactamase activity) | Alive cells | 0.29 |
|  | Heat-killed cells | 0 |
| **TEM Single mutant**<br>(moderate β-lactamase activity) | Alive cells | 0.43 |
|  | Heat-killed cells | 0 |
| **Wildtype**<br>(no β-lactamase activity) | Alive cells | 0 |
|  | Heat-killed cells | 0 |

### 3.4 Modeling the dynamic interactions between resistant and susceptible strains

Motivated by the observation that only strains expressing TEM β-lactamase reduce the concentration of CTX, we attempted to predict the amount of CTX removal from the agar medium in the experiment described above (Figure 2). In related work (Saebelfeld et al., bioRxiv preprint 2021), we obtained an estimate of the establishment probability of the Triple mutant grown alone on agar under similar conditions as a continuous function of CTX concentration through curve-fitting on a larger data set (Supplement 2: Figure S1). If we assume that the increased establishment probability in the presence of other cells in the experiment presented in Figure 2 is solely due to CTX breakdown, we can use the establishment probabilities to estimate the CTX reduction by finding the point on the curve corresponding to the increased establishment probability and its corresponding concentration (cf. Supplement 2: Figure S1). Using this, we estimate that the TEM-1 Ancestor reduces the concentration by about 0.06 μg/ml at 1.44 μg/ml CTX (4% reduction), and by about 0.25 μg/ml at 1.76 μg/ml CTX (14% reduction), whereas the Single mutant reduces the concentration by about 0.21 μg/ml at 1.44 μg/ml CTX (15% reduction), and by about 0.39 μg/ml at 1.76 μg/ml CTX (22% reduction). These estimates are subject to large uncertainties but appear nevertheless to be roughly consistent with the results of the bioassay. However, our estimates also indicate that the CTX reduction by the Ancestral strain is surprisingly high, considering its approximate 130-fold lower catalytic efficiency relative to the Single mutant.

To explain the unexpectedly high antibiotic removal by the Ancestor, we used a modelling approach. We begin with considering the difference in catalytic efficiency (*k_cat_/K_M_*) of the Ancestor and Single mutant TEM variants. Based on reported estimates for both variants (Salverda et al., 2011) and assuming constant conditions, we predict a net reduction of CTX from the medium of 0.06% for the TEM-1 Ancestor and 7.5% for the Single mutant (Supplement 3). While the prediction for the Single mutant can be considered roughly consistent with our experimental finding (given the crude nature of the estimate and the data), the prediction for the Ancestor is two orders of magnitude smaller and inconsistent with its observed only one-third smaller reduction (Table 2). Indeed, given that TEM-1 has very low activity, it is surprising that it achieves even a measurable reduction. One possible explanation is that the ancestral enzyme (TEM-1), while breaking down the antibiotics at a very low rate, nonetheless binds the antibiotic at a high rate (Queenan et al., 2004). However, this explanation seems unlikely, because removal by binding happens only for one antibiotic molecule per enzyme molecule in the periplasmic space, whereas the turnover rate due to degradation by the Single mutant is about 810 antibiotic molecules per enzyme molecule in 30 minutes (which is the scale of cell division time; the number of antibiotic molecules degraded is obtained from the reaction rate estimates in Supplement 3). Thus, binding without hydrolysis cannot explain the finding that the Ancestor and Single mutant remove CTX at comparable rates.

To investigate this issue further, we used a model introduced in Geyrhofer and Brenner (2020) which incorporates not just enzyme kinetics but also cell growth and death, and the exchange of enzyme and antibiotics between cells and the environment. Details on the dynamical model can be found in Supplement 4. In short, this model presents a more realistic picture of the dynamical process underlying antibiotic removal. Cell numbers decrease initially at high antibiotic concentration, while simultaneously removing antibiotic through β-lactamase synthesis and secretion. Depending on the inoculum size, the decrease in antibiotic concentration in the medium can lead to a resurgence in cell growth at a later time (see Figure 3). The model has a multi-dimensional parameter space that can be explored to locate the source of the unexpectedly high antibiotic removal by the Ancestral strain. Many of the parameters in the model can be either determined from the literature or have an insignificant impact on the results (Supplement 4). However, two important parameters that are left undetermined are *σ_B_* and *σ_E_*, which are the rate constants for the flux of antibiotic and enzyme, respectively, between the cells and the environment.

**Figure 3:**
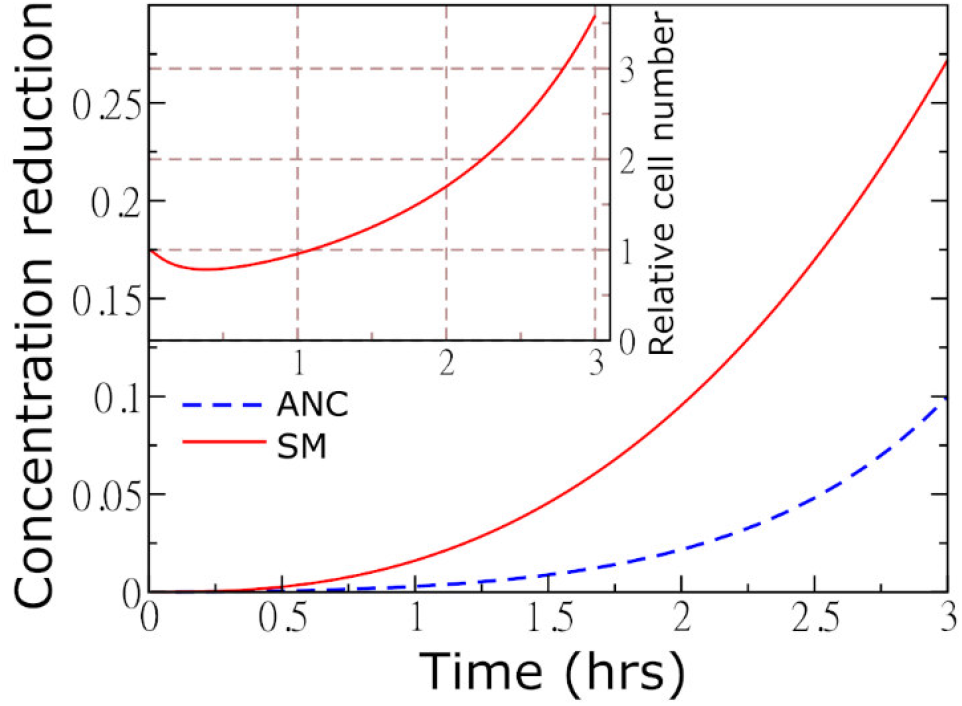
The reduction of concentration as a function of time. Parameters: TEM-1 Ancestor (ANC) *σ_E_* = 16 *h*^-1^; *σ*_B_ = 9.2 *h*^-1^; single mutant *σ_E_* = 2.81 *h*^-1^; *σ_B_* = 1 *h*^-1^. The inset shows the number of Single mutant (SM) cells relative to the initial number as a function of time. The cells die initially and then the numbers grow again as antibiotic concentration is reduced. The Ancestor cells are assumed to filament but not to divide or die.

One possibility is that the enzyme synthesis and/or permeability parameters *σ_B_* and *σ_E_* differ for TEM-1 Ancestor and Single mutant. For example, *σ_B_* and *σ_E_* could be higher in the TEM-1 Ancestor due to higher rates of cell wall defects, although there is no empirical support for this to our knowledge. However, numerical simulations show that the discrepancy is not explained even when *σ_E_* and *σ_B_* are two orders of magnitude higher for the Ancestor. As this phenomenon by itself does not provide a satisfactory explanation, we considered a possible additional role of cell filamentation. We know that our *E. coli* strain undergoes filamentation under CTX exposure, and both the rate of filamentation (Jeffrey Power, personal communication) and time of subsequent cell lysis (Zahir et al., 2020) are strongly strain-dependent. Further, during filamentation, the biomass of the cells keeps growing, even though cell division does not occur due to inhibited septum formation (Pogliano et al., 1997). Filamenting cells, therefore, continue to synthesize enzyme that breaks down the antibiotic, even if many of them eventually lyse and die (Zahir et al., 2020). If we assume that in our experiments the TEM-1 Ancestor cells do filament (and do not lyse within the experimental duration) and that enzyme production is proportional to biomass growth whereas this is not the case for the Single mutant, this can be incorporated in the model by setting the growth rate parameter to be independent of the antibiotic concentration. Under this assumption, the breakdown by the Ancestor becomes comparable to that of the Single mutant in the relevant parameter range. For *σ_E_* = 2.81 *h*^-1^, *σ_B_* = 1 *h*^-1^, the reduction is 1.8%, which is significantly higher than the value obtained without filamentation, but still lower than what is expected from the bioassay data (see Supplement 4: Figure S2). But moderately higher values of enzyme expression and release produce better results. For example, with *σ*_E_ = 16 *h*^-1^, *σ_B_* = 9.2 *h*^-1^ we get a 10% reduction. The simulated reduction of antibiotic concentration by filamenting ancestral cells is shown in Figure 3.

## 4 Discussion

We examined whether and how the establishment of *de novo* antibiotic-resistant mutant cells was affected by more susceptible resident cells. The successful establishment of a mutant cell was determined by its outgrowth into a colony-forming unit, visible to the naked eye. We first showed that the establishment probability of three separately tested TEM β-lactamase mutants with increasing hydrolysis activity against the cephalosporin CTX was increased when introduced into populations of the Ancestor strain with very low β-lactamase activity under exposure to two CTX concentrations. This positive effect increased with the density of Ancestor cells.

We next hypothesized that three, mutually not exclusive, mechanisms could be involved in enhancing mutant establishment probability. First, as the Ancestor cells died in the presence of the antibiotic, additional nutrients provided by cell lysis could support mutant establishment. It has been shown repeatedly that contents released from decaying bacterial cells can be reutilised by growing cells (e.g. Koch, 1959; Banks and Bryers, 1990; Rozen et al., 2009). Coates et al. (2018) found that the establishment of antibiotic-resistant *E. coli* mutants under antibiotic exposure was lower in a poor than in a rich nutrition environment due to a reduction in growth rate. As we used M9 minimal medium in our experiments, it is conceivable that extra nutrients released from dying Ancestor cells supported mutant establishment. Second, the Ancestor cells may have depleted CTX molecules via binding to penicillin-binding proteins (PBPs), which are the main target of CTX (Cho et al., 2014), thereby reducing the antibiotic concentration in the medium (Udekwu et al., 2009; Abel Zur Wiesch et al., 2015). Third, Ancestor cells may have reduced the environmental CTX concentration by catalyzing its hydrolysis, even though the β-lactamase activity against CTX of this strain is low.

A positive effect of susceptible cells on the establishment of *de novo* antibiotic-resistance conferring mutations has been reported before for *P. aeruginosa* in the presence of streptomycin (Alexander and MacLean, 2020). The authors speculated that the mechanism for this effect was the binding of the antibiotics to components of the susceptible cells. The main target for streptomycin is ribosomes (Demirci et al., 2013), a major cell component with numbers ranging between a few thousand to several ten-thousands per cell, depending on growth rate and species (Martin and Iandolo, 1975; Bremer and Dennis, 1996; Barrera and Pan, 2004; Leskelä et al., 2005). CTX predominantly binds to PBP3 (Jones and Thornsberry, 1982; Piras et al., 1990), which is present at about 60 to 130 units per *E. coli* cell, depending on the medium and growth rate (Dougherty et al., 1996). Thus, the antibiotic reduction via binding to cellular components can be expected to be much lower in our experimental system compared to that of Alexander and MacLean (2020). Indeed, we found that the Wildtype strain without β-lactamase was not able to remove detectable amounts of CTX from the liquid medium (Table 2). This indicates that the removal of CTX via binding to cellular components is negligible in our system.

To test whether nutrient release from decaying cells promoted mutant establishment, we used heat-killed TEM-1 Ancestor and Single mutant cells. Heat-killing was chosen to inactivate the β-lactamases as hydrolyzing enzymes can further break down antibiotics after cell death (Udekwu et al., 2009; Lenhard and Bulman, 2019). To sustain one cell, an estimated 110 heat-killed *E. coli* cells are needed to provide sufficient nutrients (Nioh and Furusaka, 1968). Thus, some 16,500 heat-killed Ancestor or Single mutant cells would have been needed to sustain the growth of the 150 plated Triple mutant cells under starving conditions. The absence of an effect of 10^6^ added heat-killed cells on the establishment of the Triple mutant (Figure 2) suggests that the addition of nutrients due to cells lysis plays a negligible role under the tested conditions, leaving CTX removal from the environment as the most probable explanation for the observed positive effect on mutant establishment. However, we cannot rule out that our method of heat-killing causes only a fraction of the cells to lyse and release nutrients (Smakman and Hall, bioRxiv preprint 2020).

The establishment of the Triple mutant was increased when introduced into the moderately-active Single mutant population compared to the low-activity TEM-1 Ancestor (see Figure 2), indicating that hydrolysis of the antibiotic contributes to the observed effect. This was further supported by the results of the bioassay, where both the TEM-1 Ancestor and the TEM Single mutant were able to remove CTX from the liquid medium, whereas the Single mutant removed more of the antibiotic than the Ancestor (Table 2). However, the CTX removal rate by the Ancestor and its effect on Triple mutant establishment are unexpectedly high when compared with the Single mutant. Specifically, the Ancestor removed about two-thirds of the amount of CTX (Table 2) and restored roughly half of the lost established Triple mutant cells compared to the Single mutant (Figure 2), despite its 130-fold lower catalytic efficiency against CTX (Salverda et al., 2011). Thus, CTX hydrolysis based on enzyme kinetics alone cannot explain the observed effect of the Ancestor. To investigate this further, we adapted a model introduced by Geyrhofer and Brenner (2020), where the effect of cell filamentation could be incorporated by allowing the biomass of cells to grow without cell division in the presence of the antibiotic. Exploration of various scenarios using this model indicates that delayed lysis of filamentous cells, coupled with an increased β-lactamase release due to cell wall permeability and increased β-lactamase synthesis rate, may explain the unexpectedly high breakdown of CTX by the TEM-1 Ancestor. While these results are not conclusive, they do point to interesting directions for future work.

Previous studies comparing antibiotic resistance evolution on agar and liquid medium showed strong differences regarding various genotypic and phenotypic outcomes. Specifically, growth on agar generally leads to a higher diversity due to greater heterogeneity in environmental conditions and inefficient competition (Rainey and Travisano, 1998; Kerr et al., 2002; Habets et al., 2007; Frost et al., 2018; Kayser et al., 2019). In related work (Saebelfeld et al., bioRxiv preprint 2021), we show that structured environments can affect mutant establishment due to greater variation in local conditions. These findings may have clinical implications as agar conditions better resemble natural conditions than liquid medium since most bacteria grow in biofilm or attached to surfaces (Olsen, 2015; Gupta et al., 2016; Ahmed et al., 2018). The maintenance of resistant cells within sensitive populations in the absence of antibiotics leads to a higher risk of colonization by the mutant when antibiotics are applied (De Roode et al., 2004; Pollitt et al., 2014; Olsen, 2015). Here, we show that this emergence could be further supported when the antibiotic is broken down. Apart from hydrolyzing enzymes like β-lactamases, those systems could include other antibiotics that are modified or degraded, such as erythromycin, tetracyclines and chloramphenicol (Nicoloff and Andersson, 2016).

In conclusion, we showed that the removal of CTX from the environment by bacterial cells expressing β-lactamase enzymes with relatively low activity against the antibiotic enhanced the establishment probability of mutants with more active enzymes in a structured environment. We believe that similar positive effects may affect the establishment of mutants resistant to other antibiotics, as was shown for streptomycin (Alexander and MacLean, 2020), but the size of this effect will depend on system-specific factors, including the antibiotic target and resistance mechanism, the ability of the strain for filamentation and the environmental structure. Notably, the positive effect of susceptible cells on the initial establishment of resistant mutants we report contrasts with their later impeding influence by preventing the fixation of higher resistance mutants once sufficient antibiotic has been removed (Yurtsev et al., 2013). Our study is among few demonstrating the importance of the resistance mechanism and conditions during the emergence of antibiotic resistance. Further investigations on factors contributing to the establishment of *de novo* resistance mutants in other systems would contribute to a better understanding of the long-term persistence and re-emergence of infections with resistant pathogens in clinical settings.

## 5 Conflict of Interest

The authors declare that the research was conducted in the absence of any commercial or financial relationships that could be construed as a potential conflict of interest.

## 6 Author Contributions

All authors conceived the study. MS, JB and AH conducted the experiments. SGD developed the model with help from JK. MS and SGD wrote the manuscript with help from JK and JAGMdV. All authors read the manuscript and approved the final version.

## 7 Funding

This work was supported by the German Research Foundation (DFG; CRC 1310 Predictability in Evolution).

## 8 Acknowledgements

We would like to thank Jeffrey Power for helpful discussions on the manuscript and insight into his data.

## Notes

### Competing Interest Statement

The authors have declared no competing interest.

