## Supplementary material for "Antibiotic breakdown by susceptible bacteria enhances the establishment of β-lactam resistant mutants"

### 1 ANOVAs and posthoc tests of the Interaction experiments

#### 1.1 Interaction experiment 1

**Table S1:** Effect of TEM-1 Ancestor presence on the establishment of three tested TEM mutant strains presented in Figure 1 (main article). Results of the two-way ANOVAs run per strain for effects of added TEM-1 Ancestor cells (*anc*), CTX concentration (*ctx*) and Ancestor cells by CTX interaction (*anc:ctx*), and results of Tukey's HSD tests. F-statistics of the ANOVAs are shown across all treatment comparisons per strain; results of Tukey's HSD tests are shown only for comparisons between the number of added Ancestor cells within each CTX concentration (per strain). Significant p-values are depicted in bold. All data were *sqrt* transformed. *N* = 15, except for the 0.14 µg/ml CTX (Single mutant) treatment where *n* = 14 per tested number of Ancestor cells.

| **Two-way ANOVA** | **Strain** | **CTX concentration (µg/ml)** | **Number of Ancestor cells** | | |
|---|---|---|---|---|---|
| | | | *0 vs. $10^3$* | *0 vs. $10^6$* | *$10^3$ vs. $10^6$* |
| *anc* $F_{2,149}$=100, **p<0.001**<br>*ctx* $F_{1,588}$=789, **p<0.001**<br>*anc:ctx* $F_{2,23}$=15.6, **p<0.001** | Single mutant | 0.14 | **<0.001** | **<0.001** | 0.65 |
| | | 0.20 | **<0.05** | **<0.001** | **<0.001** |
| *anc* $F_{2,25}$=16.3, **p<0.001**<br>*ctx* $F_{1,25}$=32.2, **p<0.001**<br>*anc:ctx* $F_{2,2}$=1.2, p=0.30 | Double mutant | 1.12 | 0.72 | **<0.001** | **<0.05** |
| | | 1.28 | 0.99 | 0.08 | **<0.05** |
| *anc* $F_{2,18}$=16.2, **p<0.001**<br>*ctx* $F_{1,19}$=33.0, **p<0.001**<br>*anc:ctx* $F_{2,0.23}$=0.2, p=0.81 | Triple mutant | 1.44 | 0.78 | **<0.01** | 0.15 |
| | | 1.76 | 0.24 | **<0.001** | 0.34 |

### 1.2 Interaction experiment 2

**Table S2:** Effect of alive or heat-killed TEM-1 Ancestor or TEM Single mutant presence on the establishment of the Triple mutant. Results of a two-way ANOVA for effects of added background population (*pop*; cf. Figure 2 in the main text), CTX concentration (*ctx*) and background population by CTX interaction (*pop:ctx*), and results of the posthoc Tukey's HSD test. F-statistics of the ANOVA are shown across all treatment comparisons; results of Tukey's HSD test are shown only for comparisons between background populations within CTX concentrations. Significant p-values are depicted in bold; *n* = 15.

| **Two-way ANOVA** | **Strain/population combinations** | **CTX concentration (µg/ml)** | |
|---|---|---|---|
| | | *1.44* | *1.76* |
| *pop* $F_{4,40375}$=425.7, **p<0.001** *ctx* $F_{2,55998}$=1180.8, **p<0.001** *pop:ctx* $F_{4,553}$=5.84, **p<0.001** | *Triple mutant vs. heat-killed Ancestor* | 0.99 | 0.92 |
| | *Triple mutant vs. heat-killed Single mutant* | 0.99 | 0.99 |
| | *Triple mutant vs. alive Ancestor* | **<0.001** | **<0.001** |
| | *Triple mutant vs. alive Single mutant* | **<0.001** | **<0.001** |
| | *Heat-killed Ancestor vs. heat-killed Single mutant* | 0.94 | 0.99 |
| | *Heat-killed Ancestor vs. alive Ancestor* | **<0.001** | **<0.001** |
| | *Heat-killed Ancestor vs. alive Single mutant* | **<0.001** | **<0.001** |
| | *Heat-killed Single mutant vs. alive Ancestor* | **<0.001** | **<0.001** |
| | *Heat-killed Single mutant vs. alive Single mutant* | **<0.001** | **<0.001** |
| | *Alive Ancestor vs. alive Single mutant* | **<0.001** | **<0.001** |

### 2 Determining establishment probability and the corresponding concentration

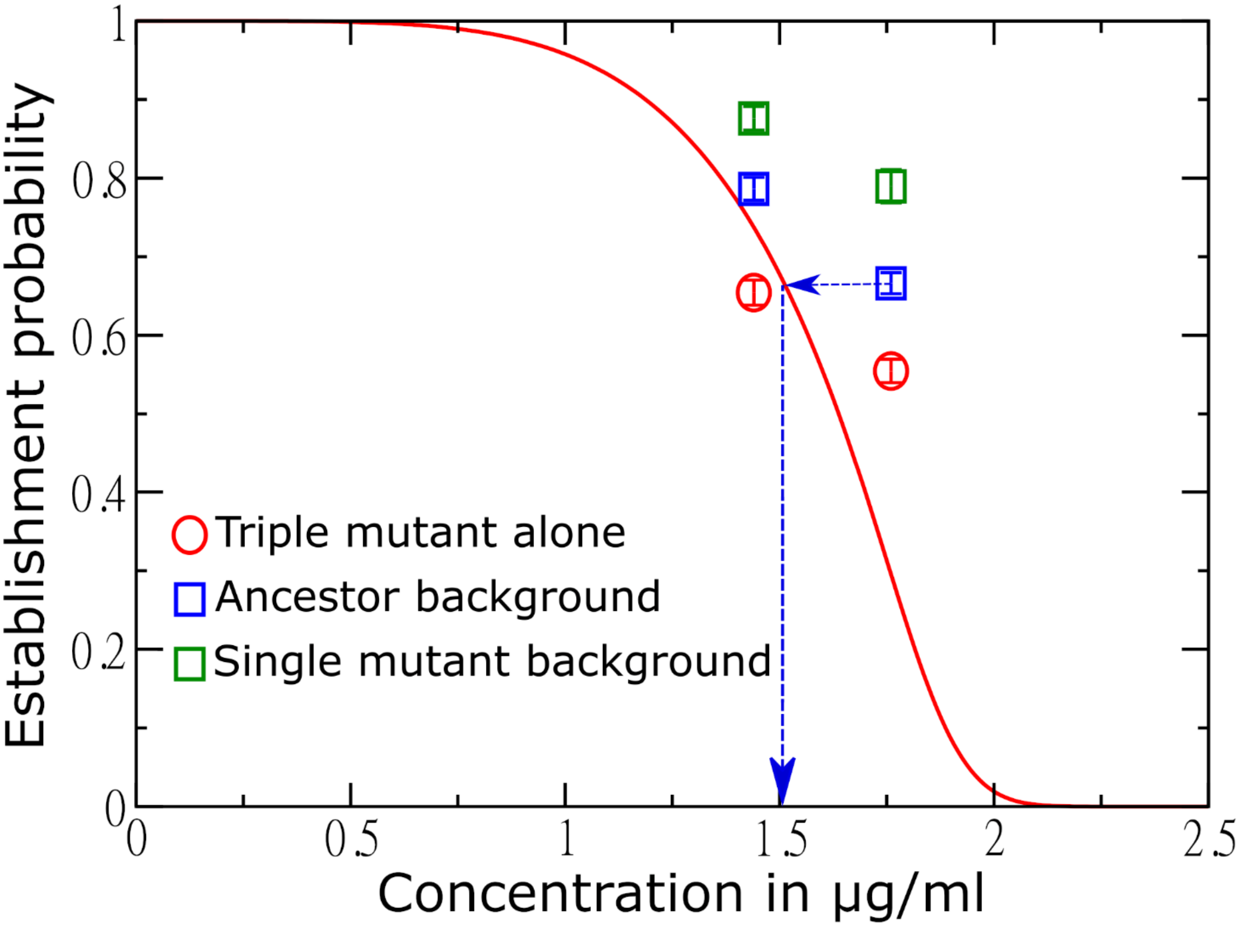


**Figure S1:** Inferred establishment probability of the Triple mutant in the absence and presence of Ancestor and Single mutant background cells. The establishment probabilities were obtained as the ratio of the mean CFU in the presence of 1.44 or 1.76 μg/ml CTX to that in the absence. The solid curve is obtained from a different study performed under the same conditions (Saebelfeld et al., bioRxiv preprint 2021), but with a larger set of CTX concentrations. The curve was determined from a model that assumes a simple branching process, where the cell division probability was taken to have a Hill function form $p_d = \frac{1}{1+\left(\frac{x}{x_0}\right)^n}$ and the parameter $x_0$ is a random variable that varies across cell lineages according to a gamma distribution. Curve fitting was performed to obtain n = 5.01 and a mean of 1.89 μg/ml and a standard deviation of 0.102 μg/ml for $x_0$. The estimate of the decreased concentration is done using the solid curve; as an example, one may determine the new, reduced concentration in the presence of the TEM-1 Ancestor at 1.76 μg/ml by following the blue arrows. The discrepancy between the red symbols and the full curve reflects experimental uncertainties and inherent limitations of the model and provides a rough measure of the accuracy of the inferred concentration changes.

### 3 Expected CTX reduction in the bioassay

To understand why the CTX reduction in the bioassay by the TEM-1 Ancestor compared to the Single mutant is unexpectedly high, we begin with a simple calculation based on chemical kinetics. Let the concentration of antibiotic in the periplasmic space be $B_{in}$, and the enzyme concentration there be $E$. Then, the rate of reduction of concentration in the cell is

$$-\frac{dB_{in}}{dt} = \frac{k_{cat}}{k_M} E B_{in}. \tag{1}$$

For the Single mutant, we have $k_{cat}/k_M = 1.3 \times 10^5\ s^{-1}\ M^{-1}$ (Salverda et al., 2011). The enzyme concentration is 168 molecules per cell (Suvorov et al. 2007 found that about 12,600 molecules are expressed from about 75 plasmid copies, leading to the estimate of 168 molecules per gene copy). A cell volume is taken to be about 1 μm$^3$ (Kubitschek, 1969; Wang et al., 2008), and the periplasmic space volume is 0.3 μm$^3$ (Stock et al., 1977). Therefore the enzyme concentration is $E = 0.93$ μM. If the antibiotic concentration within the periplasmic space is the same as the initial concentration in the medium, 1.6 μg/ml (which is the average of the two different values used in the bioassay) or 3.6 μM (using the molecular weight 445 g/mol and the volume of periplasmic space); then the net reduction rate in the periplasmic space is computed from above to be 0.19 μg/ml per second per cell or 76 molecules per second per cell. Assuming a constant number of 2x10$^8$ cells and 3 hours incubation, one predicts a net reduction of 0.12 μg/ml in the medium, i.e. 7.5%. Similarly, for the TEM-1 Ancestor, $k_{cat}/k_M = 10^3\ s^{-1}$ (Salverda et al., 2011), and the reduction is predicted to be $9.2 \times 10^{-4}$ μg/ml or 0.06%.

### 4 Dynamical model

We adopt a dynamical model from Geyrhofer and Brenner (2020). We define $N$ as the number of cells in the medium, $B_{in}$and $B_{out}$ as the antibiotic concentration inside and outside the cell respectively, and $E_{in}$ and $E_{out}$ as the concentration of enzyme inside and outside the cell, respectively. Then the system is described by the following equations

$$\frac{dN}{dt} = \alpha_e(B_{in})N$$

$$\frac{dB_{in}}{dt} = -\varepsilon E_{in} B_{in} + \sigma_B(B_{out} - B_{in})$$

$$\frac{dB}{dt} = \eta\sigma_B N(B_{in} - B_{out}) - \varepsilon E_{out} B_{out} \tag{2}$$

$$\frac{dE_{in}}{dt} = \rho + \sigma_E(E_{out} - E_{in}) \tag{3}$$

$$\frac{dE_{out}}{dt} = \eta N \sigma_E(E_{in} - E_{out}),$$

where we take the growth rate $\alpha_e$ to have the empirical form (Geyrhofer and Brenner, 2020)

$$\alpha_e(B_{in}) = \alpha \frac{1-\left(\frac{B_{in}}{\mu}\right)^{\kappa}}{1+\frac{1}{\gamma}\left(\frac{B_{in}}{\mu}\right)^{\kappa}}. \tag{4}$$

The terms containing $\sigma_B$ and $\sigma_E$ correspond respectively to the fluxes of antibiotic and enzyme between a cell and the medium, and the quadratic terms containing $\varepsilon$ are the breakdown rates of antibiotic by the enzyme. Note that the breakdown is taken to occur both within and outside the cell. We take $\gamma = 1$ (Geyrhofer and Brenner, 2020) and $\kappa = 2$ (the precise value does not matter); $\alpha$ is the growth rate in the absence of antibiotics, which we take to correspond to a division time of 30 minutes. The parameter $\mu$ is the MIC of a cell in the absence of any enzyme activity, such as the cells of the Wildtype strain without the TEM allele in the bioassay. We do not know this value, but we can establish upper and lower bounds and estimate the single-cell MIC as the geometric mean of the two. It should be less than the MIC of the TEM-1 Ancestor, which is 0.04 µg/ml (based on single-cell MICs run in triplicates with 2-fold increases in CTX concentration and 4-fold decreases in inoculum size starting from $10^5$ cells; data not shown), or about 52 molecules in 1 $\mu m^3$, which is the cell volume. We do not expect the MIC to be less than 1 molecule per cell, establishing the lower bound. The geometric mean gives us the estimated MIC, $\mu = 7.2$ molecules per $\mu m^3$. The volume separation factor η is the ratio of the volume of the periplasmic space (about 30% of the cell volume (Stock et al., 1977), which is 1 $\mu m^3$) to the volume of the medium, which is 1 ml. This comes out as $\eta = 3 \times 10^{-13}$. The reaction rate constant $\varepsilon = \frac{k_{cat}}{k_M}$ is $0.78\ (\mu m^3 hr)^{-1}$for the Single mutant, and $6.0 \times 10^{-3}\ (\mu m^3 hr)^{-1}$ for the TEM-1 Ancestor (Salverda et al., 2011). The synthesis rate of the enzyme is ρ (Eq. 3), whereas $\sigma_E$ is the rate constant for enzyme secretion from the cells. When the enzyme concentration outside the cell is low, the steady-state concentration of the enzyme in the cells is given $E_{in}^{ss} = \frac{\rho}{\sigma_E}$. We use the value of 168 molecules per cell (see the previous subsection) for $E_{in}^{ss}$ , and this fixes the ratio of $\rho$ and $\sigma_E$. Determining one of them fixes the other. The rate constant for antibiotic flux through the cell is $\sigma_B$.

We are finally left with two undetermined parameters, $\sigma_B$ and $\sigma_E$, and we need to explore this parameter space to understand the amount of reduction achieved. Figure S2 shows the simulation results for this model, over large ranges of the two parameters $\sigma_B$ and $\sigma_E$. The solid lines belong to the Single mutant. The parameter regime we are interested in should produce a breakdown of a few tens of per cent, and growth in cell number by a factor of about 10 or less (corresponding to a generation time of about 1 hour and a similar lag time preceding growth; based on previous observations with the tested strains under similar conditions, data not shown). By these criteria, the relevant ranges are seen to be $\sigma_B$ about $1 \sim 3\ h^{-1}$, and $\sigma_E$ in the range $1 \sim 10\ h^{-1}$. (A more detailed numerical exploration shows that the values $\sigma_E = 2.81\ h^{-1}$; $\sigma_B = 1\ h^{-1}$ produce a reduction of approximately 27% and a final cell population 3.6 times higher than the initial value, close to the value 4 we would get with one hour lag and one hour doubling time). The precise parameter and reduction values are unimportant; we are interested in whether the Ancestor cells have the same order-of-magnitude reduction in the relevant parameter range. The dashed lines in Figure S2 show the reduction of antibiotic by the Ancestor, and the reduction is about 2 to 3 orders of magnitude lower than that due to the Single mutant in our region of interest. Specifically, for $\sigma_E = 2.81\ h^{-1}$; $\sigma_B = 1\ h^{-1}$, the reduction is 0.062%, which is far too small. The experimental observation is therefore not fully explained. However, the blue dashed-dotted line includes the effect of 100% filamentation among the Ancestor cells, and it exhibits a much higher reduction - close to the actual amount of reduction observed. We implement the filamentation in Ancestor cells by setting $\alpha_e(B_{in}) = \alpha$.

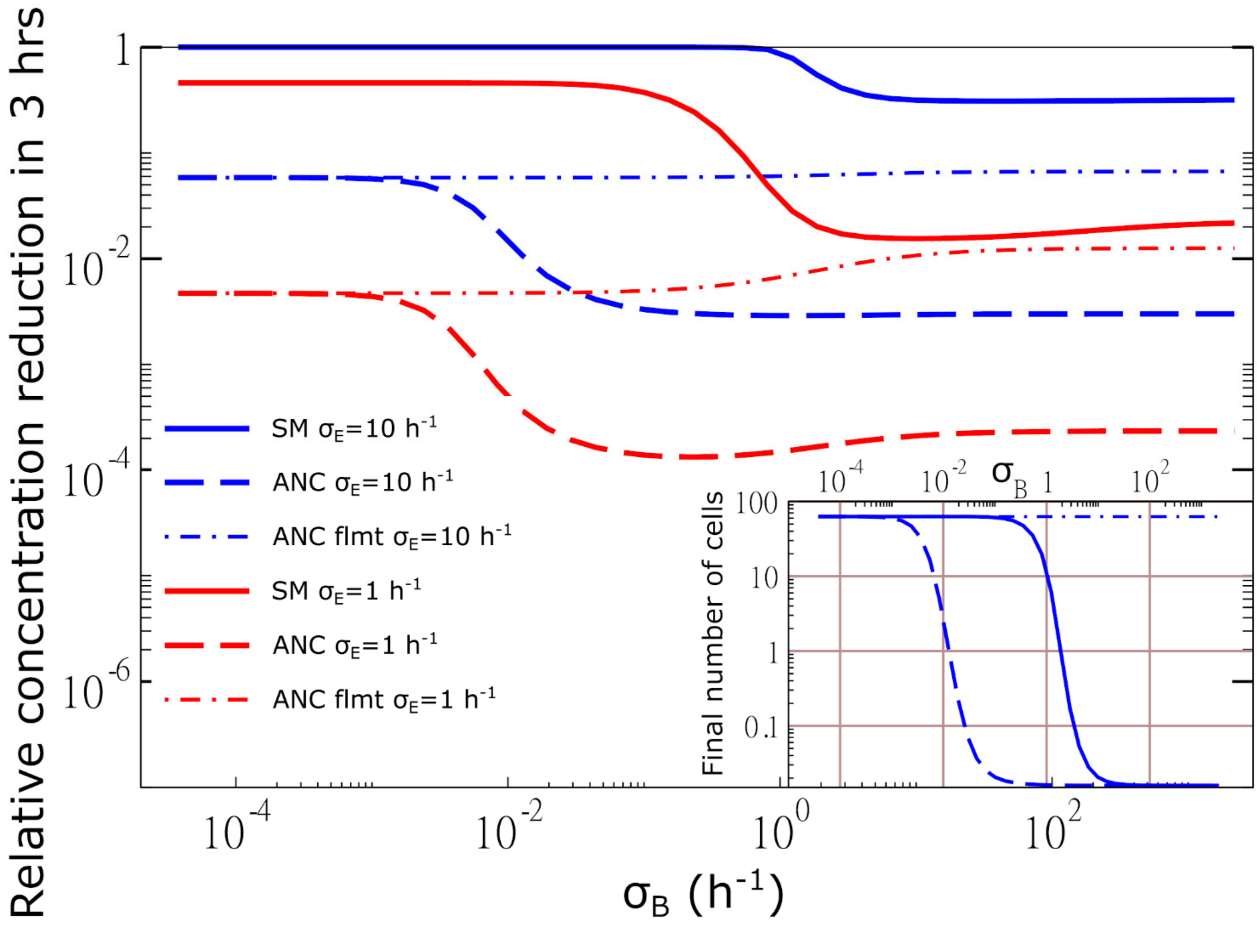


**Figure S2:** Simulation results for antibiotic concentration reduction in liquid. Solid lines are for the TEM Single mutant (SM), dashed lines for TEM-1 Ancestor (ANC) without filamentation. The dash-dotted lines are for the Ancestor with filamentation and two different enzyme transport rates. The inset shows the number of cells after 3 hours.
